# Analysis of the interactions between physiological substances and endogenous hormones in *Bombax ceiba* bud differentiation

**DOI:** 10.1101/2025.01.15.633137

**Authors:** Shuyu Wang, Zeping Cai, Ying Zhao, Deqiao Cui, Meixing Wang, Yanting Zhang, Xudong Yu

## Abstract

2

Plant bud differentiation is a process of transition from nutrient growth to reproductive growth, which is important for improving crop yield, breeding of good seeds and regulation of flowering time. To elucidate the morpho-physiological changes during bud differentiation in *Bombax ceiba*, this study was conducted in Haikou City, Hainan Province, employing paraffin sectioning, plant physiological assays, and LC-MS. We aimed to determine the dynamic changes occurring during this process. The results showed that 1) firstly, the transformation from growth point to flower bud is called floral primordial development, and secondly, the floral organ is mature.2) The content of soluble sugar and soluble starch of early-flowering and late-flowering *Bombax ceiba* first increased and then decreased, and the content of sucrose, soluble protein and malondialdehyde first decreased and then increased. The content of chlorophyll a gradually decreased and that of chlorophyll b and carotenoids gradually increased in early-flowering *Bombax ceiba*, while the opposite was true in late-flowering *Bombax ceiba*.3) 5Ds, CK and Auxin mainly acted in stage I and II, and ABA, SA, GA, ETH, and JA mainly acted in stage II and III.4) Soluble sugar and GA, soluble protein and IAA\CK, sucrose\malondialdehyde and IAA\CK\JA, soluble starch and IAA\CK\GA were significantly positively correlated, and chlorophyll and IAA\CK\SA were significantly negatively correlated, with the existence of synergistic and antagonistic effects co-regulating bud differentiation.This study provides a theoretical basis for exploring the mechanism of flower formation in *Bombax ceiba* and production practice.

**Summary statement:** To investigate the morphological and physiological characteristics of the bud differentiation process in *Bombax ceiba*, and to reveal the mechanisms of physiological substances and hormone interactions.

## 4 Introduction

Plant flowering mechanism is a complex and precise regulatory network, in which physiological perception system and hormone regulation system play an indispensable core role (Wang Rong, 2007), which accurately regulate the developmental process of floral organs by participating in the spatiotemporal specific expression of flower development. Its quality directly affects plant growth and development, and is of great significance in improving crop yield, breeding of good seeds, and regulation of flowering time. Flower bud differentiation in plants is a process of transition from nutrient growth to reproductive growth, which is the result of the synergistic action of internal and external factors. In recent years, new hormone-regulated receptors have been discovered and identified, and the mechanisms of various kinase cascades and secondary messenger systems have been gradually elucidated in the study of signal transduction pathways, and the study of hormone signals regulating the expression of flowering genes has also been deepened (Yongqiang Yu. et al., 2023; Wang, J. et al., 2024).

*Bombax ceiba* is a large deciduous tree of the genus *Bombax* in the family Bombacaceae, which combines multiple uses such as ornamental, medicinal, spinning, timber, and landscaping, and it is a rare species of ecological forests, landscape forests, and economic forests (Wu, J.S. and Yu, X.D.,2017). Hainan Island possesses abundant Bombax ceiba resources. A survey by Wang Jian et al. found that *Bombax ceiba* is distributed across the island, with the highest concentration in Baisha and Changjiang counties. Notably, Changjiang is known as China’s ‘Bombax ceiba hometown,’ and the Bombax ceiba planting area in the province accounts for approximately 17.71% of the total.

As the natural flowering period of *Bombax ceiba* is mainly concentrated in winter and spring, the ornamental period is shorter, the cultivation time is longer, and less basic research, and few studies on the interrelationship between physiological substances and endogenous hormones on the regulation of bud differentiation, less research on the mechanism of *Bombax ceiba* flower greatly affects the promotion and application of *Bombax ceiba*, in order to reveal the characteristics of the *Bombax ceiba* flower bud differentiation, to understand the reproductive strategy and adapt to the environment, and to investigate the energy metabolism and substance accumulation of *Bombax ceiba*, this study was carried out in the field of *Bombax ceiba* planting. In order to reveal the characteristics of *Bombax ceiba* flower bud differentiation, to understand the reproduction strategy and adaptation to the environment, and to investigate the energy metabolism and material accumulation of *Bombax ceiba*, this study took Haikou City, Hainan Province as the study area, and relied on the *Bombax ceiba* germplasm resource nursery of Hainan University and the Ministry of Education Key Laboratory of Tropical Speciality Forest Trees and Flower Genetics and Germplasm Innovation, to investigate the energy metabolism and material accumulation of *Bombax ceiba*. We used paraffin sections, plant physiological assays, LC-MS and other methods to measure the morphological and physiological characteristics of *Bombax ceiba* during the bud differentiation process, and to explore 1) morphological and structural characteristics and physiological changes during the bud differentiation process, 2) identification of 88 endogenous hormones and hormone-related differential metabolites during the bud differentiation process of *Bombax ceiba*, and 3) investigation of hormone and physiological substance interactions in the network of the bud differentiation stage. To provide theoretical basis and reference materials for later research on the mechanism of flower formation in *Bombax ceiba* and production practice.

## 5 Materials and methods

The test site is located in Hainan University, Haikou City, Hainan Province (20°3′19.38″N, 110°19′30.1″E) in 2021, selected 30 with the same genetic background, good growth condition, the same growth, tree height of 5.8-7.5 m, crown spoke of 4.2-6.5 m *Bombax ceiba* adult experimental standard 5 plants, collected annual branches or the current year on the new shoots as test materials, and every 3 d from the flower bud differentiation to the wilting process. The flower buds were collected from annual branches or new shoots of the year as test materials, and the trees were exposed to uniform light on all sides, and the whole process from flower bud differentiation to withering was observed and recorded every 3 d. Field-grown standard plants were simultaneously observed and recorded. Under normal conditions, ‘early-flowering’ *Bombax ceiba* cultivars used in this study began to develop buds at a lower height and with fewer leaves, typically 5–10 days earlier than normal-flowering *Bombax ceiba*. Conversely, ‘late-flowering’ *Bombax ceiba* cultivars initiated bud development later. Standard normal flowering *Bombax ceiba* followed typical conditions

### 1. Observation of morphological structure of bud differentiation

From 15 October 2020 to 30 March 2021, 7∼10 fresh flower buds with full buds and good development were taken every 3 d and put into self-sealing bags and marked for morphological observation.

Paraffin sectioning method Flower buds were quickly cut into 3 mm×3 mm slices with a double-sided razor blade and put into FAA fixative (38% formaldehyde, glacial acetic acid, and 70% anhydrous ethanol: 1:1:18 ratio), and three slices of the cut flower buds were completely immersed in FAA fixative for at least 24 h. Flower buds were dehydrated with different gradients of ethanol (30%, 50%, 70%, 80%, 90%, and 100%) → xyleneoxide. 100%) → xylene transparency → dipping wax → embedding → slicing → patching → dewaxing → rehydration → 1% Senka red and 1% solid green staining → washing → dehydration of one slice → drying and a series of other operations, and then placed the slices in a light microscope to observe and take pictures.

Observation by in-body microscope 20 flower buds were randomly collected from the 5th node of the new shoots every 3 d from the beginning of bud differentiation of *Bombax ceiba*, and on the day of sampling, the morphology and structure of the flower buds were observed under in-body dissecting microscope by peeling off the scales of the flower buds with bare hands and observing them with in-body microscope, counting and recording the stage of differentiation of the flower buds, and the lengths of the transverse and longitudinal stems of the flower buds were measured by using vernier calipers. When more than 75% of the flower buds were in one of the above stages of differentiation, it was determined that the sampling period had entered that stage of differentiation.

### 2 Measurement of physiological substances

From 15 October 2021 onwards, 0.5 g of fresh early-flowering cotton buds were sampled at the early stage of bud differentiation (I), the calyx/petal primordial differentiation stage (II), and the stamen/gynoecium primordial differentiation stage (III), for the determination of the contents of physiological substances (soluble sugars, sucrose, soluble starch, soluble proteins, malondialdehyde (MDA), chlorophyll a, chlorophyll b, and carotenoids) of the early-flowering and late-flowering cotton buds respectively. carotenoids) content. Three replicates were used in each group, in which soluble sugar, sucrose and soluble starch were determined by anthrone colourimetric method, soluble protein was determined by Caulophyllum blue G-250 method, malondialdehyde (MDA) content was determined by thiobarbituric acid method, and chlorophyll was determined by 95% ethanol milling and filtration method, and the methods were referred to the ‘Experimental Guidelines of Plant Physiology’.

### 3 Determination of endogenous hormones and differential metabolites

From 15 October 2021 onwards, 24 *Bombax ceiba* adult experimental standard strain buds were selected for treatment and determination of hormone levels. 0.5 g of *Bombax ceiba* buds were collected at each stage and stored at ultra-low temperature and set up three replicates, which were ground with a grinder (30 Hz, 1 min) until powdered and added to 10 μL of the internal standard mixing solution with a concentration of 100 ng/mL, 1 mL of methanol/water/formic acid (15:4:1, v/v :1, v/v/v) extractant were mixed well; vortexed for 10 min at 4 °C, centrifuged at 12000 r/min for 5 min, and the supernatant was taken to a new centrifuge tube for concentration; after that, 100 μL of 80% methanol/water solution was reconstituted over 0.22 μm filter membrane for LC-MS/MS analysis.

### 4 Data analysis

Excel software was used to categorise and organise the data, and SPSS 27.0 software was used to analyse the data by ANOVA and Duncan’s new complex polarity method for multiple comparisons, and Power point plotting.

## 6 Results

### 1 Morphological characteristics of bud differentiation

*Bombax ceiba* bud differentiation is divided into the early stage of bud differentiation (I), the calyx/petal primordium differentiation stage (II), and the stamen/pistil primordium differentiation stage (III).The new growth of *Bombax ceiba* new shoots stops in late October and gradually lignifies, with the apical growth point raised upwards, the tip of the apical tip thin, the base broadly bulging and conical, and the cytoplasm of the cells is thick and strong, with a length of the flower bud of 0.30 cm, and the width of the flower bud of 0.21 cm (Fig.1 A, Fig.1 E). In early November, the calyx basal differentiation stage (?) was entered, in which the growth cone was sunken and widened, and the top of the growth cone was gradually flattened from oblate to oblate, and both sides of the growth point produced protuberances that grew and expanded, and the base of the differentiation elongated and gradually became wider and curved inward to form the petal primordium to become the calyx primordium, and as the growth cone continued to be widened, the base of the calyx primordium was 0.65 cm in length and 0.58 cm in width (Figs.1 B and 1 F).

In early December, we entered the stage of androgynophore/gynophore protocorm differentiation (III), as the flower buds continued to differentiate and grow, the top of the growth cone began to appear concave, and multiple protruding stamen protocorms gradually formed on the inner side of the petal protocorms (Xu, Y. & Liu, J., 2024; Malakshahi Kurdestani, A & Francioli, D., 2024), and by December By mid-to late December, a distinct stamen cluster is formed and a growth cone emerges in the middle of the stamen cluster, with the new gynobasal protuberances elongating and growing rapidly to eventually cover the entire growth cone, and the flower buds are ca. 1.37 cm in length and ca. 1.01 cm in width (Figs. 1 C and 1 G) (Xiaoxiao Li & Wan, Chuan et al., 2021).

**Fig 1.**
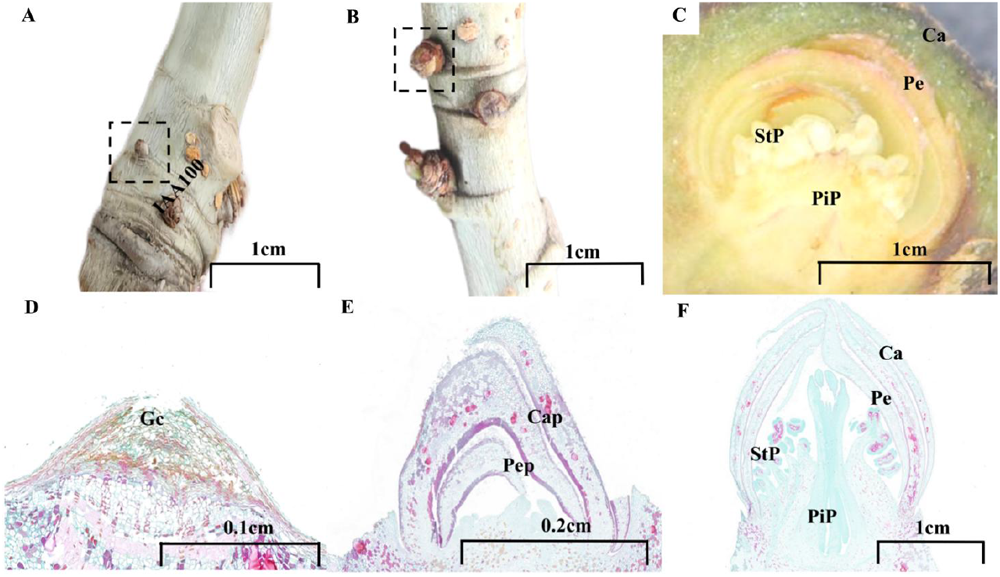
Morphological characteristics of flower bud differentiation in *Bombax ceiba*. A, E: early stage of flower bud differentiation (I); B, F: stage of calyx/petal primordium differentiation (II); C, G: stage of stamen/pistil primordium differentiation (III) Note: GC: growth cone; Cap: calyx primordium; PeP: petal primordium; Ca: calyx; Pe: petal; PiP: pistil primordium; StP: stamen primordium

After the development of the flower primordium was completed, it entered the maturation period of the floral organ under the influence of hormones in the plant and the regulation of external conditions, which was divided into five stages: the primordial stage, the fast-growing stage (3-5 d), the large balloon stage (1 d), the beginning of the flowering stage (1 d), and the full bloom stage (2-4 d). During the growing period of about 120 d, the pedicel elongates slowly, the bud elongates and enlarges slowly, and the bud becomes conical (Fig.2 A, Fig.2 B); during the fast growing period of 3-5 d, the pedicel elongates rapidly, the bud splits open at the tip, the petals are exposed and elongate rapidly (Fig.2 C); during the blooming period, the buds grow into a lo ng conic (Fig.2 D); during the initiation of flowering, most of the petals are highly revolute after they open, and a few of them stretch straight out and are thick in texture, exposing the stamen s and the pistil, and the anthers open up! Pollen dispersal, stigma secretes mucus, bees and birds collect nectar for pollination (Fig.2 E); flower buds elongate to longest and largest at ballooning stage, most petals gradually recycle inward at unopened bloom stage to form straight or included floral pattern, a few maintain original revolute floral pattern and gradually become thinner, wit h margins gradually wilting (Fig.2 F) (Chuan Wan & Dongyan Yang,2022).

**Fig 2.**
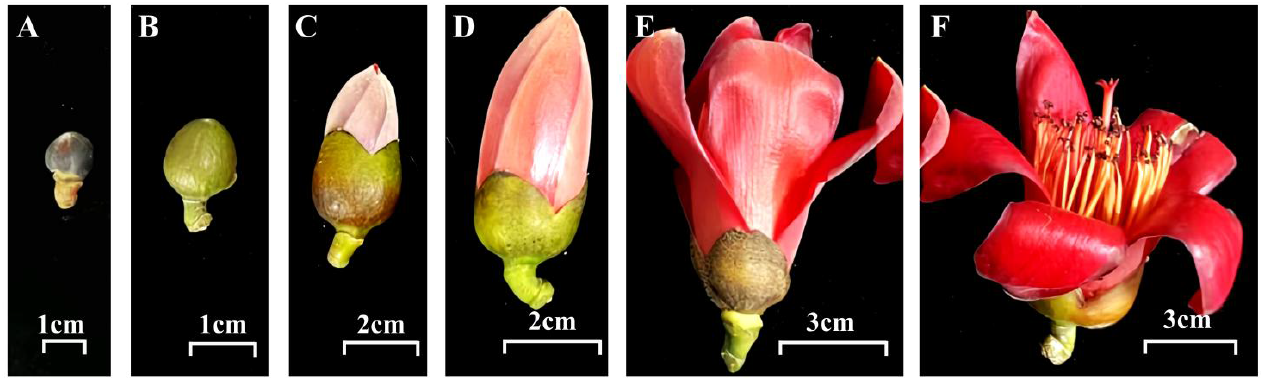
Characterization of external morphological structure of single flowers. Note:A-B: primordial stage ; C: fastigiate stage; D: large balloon stage; E: initiation stage; B-F: full bloom stage

### 2 Changes in physiological substances of bud differentiation

Physiological substances are involved in a variety of physiological processes such as energy metabolism, osmotic regulation, signalling, etc., and are an important source of energy storage in plants, as well as an important regulator of plant growth and development and gene expression (Nazir, F &. Péter, P,2024). In order to investigate the effects of flowering time and bud differentiation stage on the physiological substances of *Bombax ceiba*,two-factor ANOVA and multiple comparisons were conducted on different different differentiation stages of early-flowering and late-flowering *Bombax ceiba*, and the results showed that the soluble sugar of early-flowering *Bombax ceiba* showed a tendency to rise and then fall, and then gradually declined after rising to the peak value of 11.42 mg•g-1 in stage II, and the content of soluble sugar of late-flowering *Bombax ceiba* showed a tendency to gradually rise, and gradually rose to the peak value of 11.42 mg•g-1.The soluble sugar content of late-flowering *Bombax ceiba* showed a rising trend, gradually increasing to a peak value of 9.50 mg-g-1. in stage III, with higher significance of *0*.*06* and *0*.*43* in stages II and III, respectively (Fig. 3A). Early flowering and late flowering *Bombax ceiba* sucrose showed a decreasing and then increasing trend, the content first decreased to the lowest value of 0.80 mg•g-1. and 0.75 mg•g-1 in phase II and then gradually increased in early flowering *Bombax ceiba* more than late flowering *Bombax ceiba* content increased by 1.47 mg•g-1. and 0.78 mg•g-1. from phase II to phase III, respectively (Fig. 3A). -1. and 0.78 mg•g-1 respectively, which was most significant at *0*.*01* in phase I (Fig. 3 B). The soluble starch of early and late flowering *Bombax ceiba* showed an increasing and then decreasing trend, reaching a peak of 53.14 mg•g-1. and 47.47 mg•g-1. at stage ?, and consuming a large amount of starch after pollen mother cell development was the lowest at 12.26 mg•g-1. and 9.72 mg•g-1. with significance of *0*.*01, 0*.*01* and *0*.*03* at stages I to III, respectively (Fig. 3 C). The soluble protein content of early and late flowering *Bombax ceiba* showed a trend of decreasing and then increasing, early soluble proteins were involved in the construction of cell morphology shaping, division and differentiation to reach a minimum value of 125.08 mg•g-1,125.60 mg•g-1. at stage ?, and reached a peak value of 150.29 mg•g-1. and 174.33 mg•g-1. with a significant *0*.*01* at stage I (Fig.3 D). Malondialdehyde is an important physiological index for osmotic regulation when plants are subjected to adversity stress (Yang, D & Chen, Y, 2024; Chi, et al. 2023) The content of early-flowering and late-flowering *Bombax ceiba* showed a decreasing and then an increasing trend, gradually decreasing to 1.63 mg•g-1. in phase II, 1.68 mg•g-1. and then increased to a peak of 3.15 mg•g-1. and 2.76 mg•g-1. Early-flowering *Bombax ceiba* responded more significantly to the external temperature, which was more significant at *0*.*01* in phase III (Fig. 3 E) Chlorophylls and carotenoids in the chloroplasts during flower bud differentiation are involved in metabolism and transport and are reference indicators of plant senescence (Huang & Y.C.,2024). Early-flowering *Bombax ceiba* chlorophyll a and late-flowering *Bombax ceiba* chlorophyll b showed a tendency to gradually decrease first, increasing to peak values of 0.81 mg•g-1. and 0.26 mg•g-1. in stage I, and then decreasing to the lowest values of 0.52 mg•g-1. and 0.18 mg•g-1. Late-flowering *Bombax ceiba* chlorophyll a and early-flowering *Bombax ceiba* chlorophyll b showed a trend of gradual increase with bud differentiation, and the content of late-flowering *Bombax ceiba* chlorophyll a peaked at 0.78 mg•g-1 at stage III, with an increase of about 16%. Early flowering *Bombax ceiba* chlorophyll b content to the ? period reached a peak of 0.26 mg•g-1 after a gradual decline to 0.18 mg•g-1., chlorophyll a value of 0.26 mg•g-1. after a significant both *0*.*25* and *0*.*04* (Fig.3 F,G). Carotenoid anthocyanin deposition, petal and fruit colouration, the content of early flowering *Bombax ceiba* gradually increased from phase I to phase III, respectively, by 0.04 mg•g-1. and 0.01 mg•g-1 to a peak value of 0.27 mg•g-1. in phase III, and the content of late-flowering *Bombax ceiba* carotenoids gradually decreased to a minimum value of 0.19 mg•g-1. (Fig.3 H). Early flowering *Bombax ceiba* chlorophyll (a+b)/carotenoid and late flowering *Bombax ceiba* chlorophyll a/b content gradually decreased to peak at 4.62 mg•g-1. and after 3.82 mg•g-1. in stage I and then gradually decreased to to stage III to a minimum value of 3.02 mg•g-1. and 1.59 mg•g-1. after. Chlorophyll (a+b)/carotenoid content of late flowering *Bombax ceiba* and early flowering *Bombax ceiba* chlorophyll a/b content gradually increased and reached a peak value of 5.27 mg•g-1. and 5.21 mg•g-1. at stage III, and chlorophyll (a+b)/carotenoid was significant at all three stages (Fig. 3 I J).

**Fig 3.**
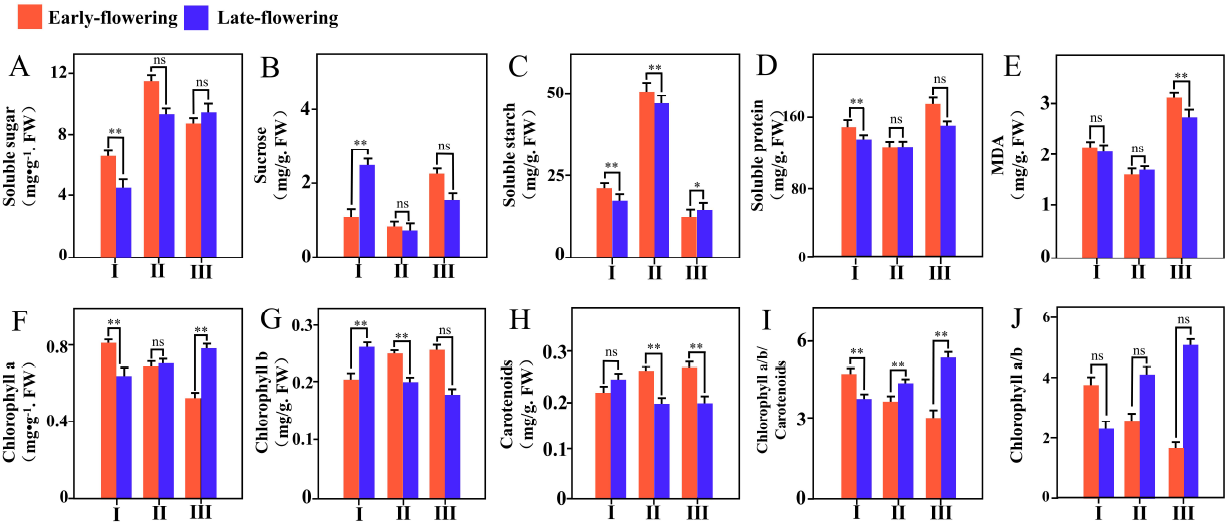
Variation of physiological substance content in flower bud differentiation. Note:A: Soluble sugar content (mg•g-1. FW); B: Sucrose content (mg•g-1. FW); C: Soluble starch content (mg•g-1. FW); D: Soluble protein content (mg•g-1. FW); E: MDA content (mg•g-1. FW); F: Chlorophyll a content (mg•g-1. FW); G: Chlorophyll b content (mg•g-1. FW); H: Carotenoids content (mg•g-1. FW); I: Chlorophyll (a+b)/Carotenoid; J: Chlorophyll a/b

### 3 Flower bud differentiation hormone levels and differential metabolites

Flowering consists of a complex flowering control network formed by multiple exogenous and endogenous signals, and phytohormones, as the most important endogenous signalling, play an important role in the process of flower formation (Zhu, Y. & Zhang, J, 2024; Huang, P.K. & Schmit, J, 2024). By determining the content of phytohormones at the stage of flower bud differentiation, 88 phytohormones existed in 8 major phytohormone categories were classified to be changed, and in general GA (GA24), 5Ds, CK (2MeSiP, BAP, iP7G, iP9G, cZ9G, oT9G, mT9G, K9G), IAA (IAA-Ala, ICA, ICAld, TRA, MEIAA, IAA-Asp) and predominantly act in phase I and II for floral primordium differentiation (Fig. 4 C E F G H). ABA (ABA, ABA-GE), SA (SA, SAG), GA (GA3, GA19), JA, IAA (MEIAA, IAA-Asp, IAA-Trp, IAN, IAA-Val-Me, IAA, TRP), CK (cZROG, 2MeScZR, DHZ7G, DHZROG, IP, IPR, DHZR, tZR, 2MeSiP), and JA (JA-ILE, MEJA, JA, JA-Val, H2JA) acted mainly in the senescence of floral organs in phase II and phase III varied significantly in phases I and II (Fig. 4 A B C F G H). ETH (ACC) content peaked at stage III (Fig. 4 D).

**Fig 4.**
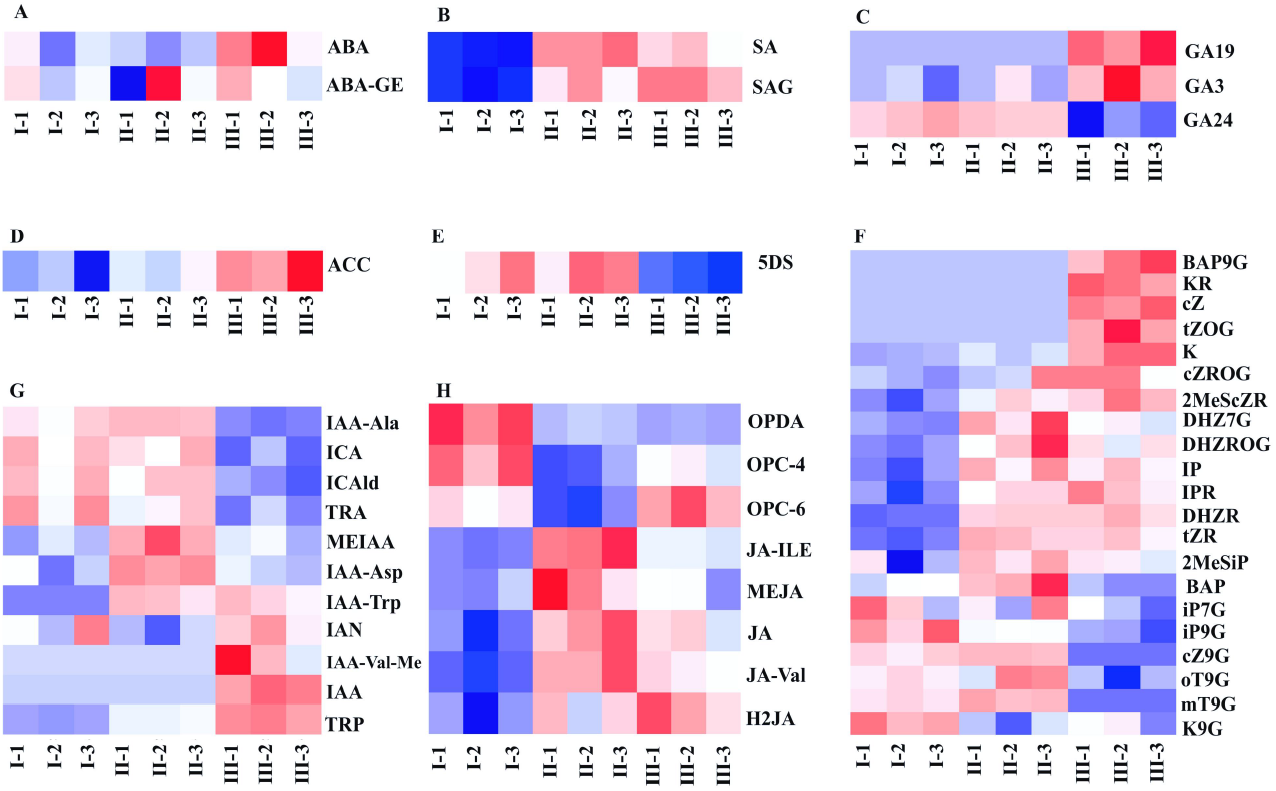
Dynamic changes of hormone levels in flower bud differentiation. Note:A:Abscisic acid analogs (ABA); B:Salicylic acid analogs (SA); C:Gibberellins (GA); D:Ethylene analogs (ETH); E:Solanum lactone (SL); F:Cytokinins (CK); G:Growth hormone (Auxin) H:Jasmonic acid (JA)

To further explore the hormone regulation-related differential metabolites during flower bud differentiation in *Bombax ceiba*, a total of 20 differential metabolites (DEMs) were screened, of which 14 were up-regulated and 6 were down-regulated, with up-regulated expression focusing on Auxin, CK, GA, and SA, and down-regulated expression focusing on Auxin, CK, JA, and 5Ds (Fig.5 A).KEGG is enriched for environmental information processing and genetic information processing, biosynthesis of secondary metabolites, metabolic pathways, phytohormone signalling on hormones and nutrients during bud differentiation are closely related to the involvement of early biological processes as response to stimuli, transmembrane translocation, cleavage enzyme activity and structural molecule activity. Zeatin biosynthesis, synthesis of various alkaloids, and metabolic pathways are associated with delayed organ senescence and storage and preservation of freshness (Fig. 5 B, C, and D). The process of bud differentiation contained a total of 24 hormone-related DEMs, with a gradual decrease in overall content, a decrease and then an increase in the content of 5Ds from stage I to stage II, and the consumption of a large amount of SA for senescence retardation by stage III (Fig.5 E). The 20 hormone-related DEMs screened were correlated, and Auxin (IAA) and CK (cZ) were strongly correlated, synergising with each other to promote cell division and floral organ growth, and growth hormone polar transport. Whereas Auxin (IAA-Val-Me) and CK (KR) antagonise each other, cytokinin promotes apical dominance and facilitates apical bud growth, and cytokinin inhibits apical dominance and promotes flower bud differentiation. (Fig. 5 F).

**Fig 5.**
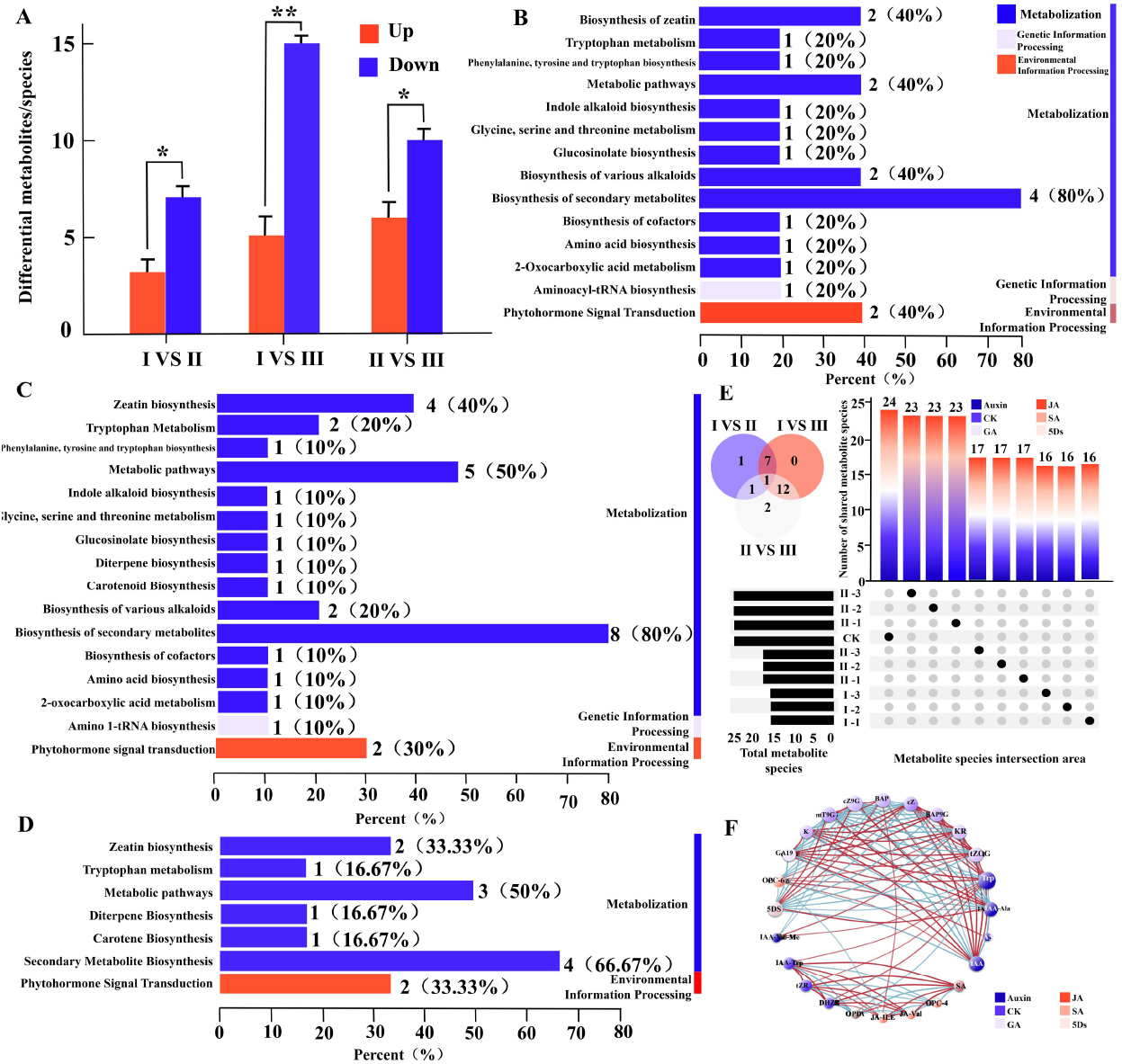
Screening of differential metabolites of flower bud differentiation. Note:A: Differential metabolite content map; B: Differential metabolite KEGG classification map (stage I and II); C: Differential metabolite KEGG classification map (stage I and III); D: Differential metabolite KEGG classification map (stage II and III); E: Differential metabolite Venn and Upset maps; F: Differential metabolite related network map

### 4 Physiological substances and endogenous hormone interaction networks

Physiological substances and endogenous hormones have mutual synergistic and resistant effects in epigenetic regulation and flowering promotion pathways. They were divided into two groups according to the degree of association, physiological substances MDA, sucrose, soluble protein and chlorophyll a were closely related with an average radius of 2.737, and the rest of them were weakly related with an average radius of 15.596. Hormones IAA, CK, and 5Ds were closely related with an average radius of 2.854, and had the highest degree of association with physiological substances Sucrose, MDA, soluble protein, and chlorophyll a, whereas JA, SA had lower correlation (Fig 6-B).Mantel test analysis and clustering heat map showed significant positive correlation between soluble sugars and G19 *(0*.*01<p<0*.*05)*, soluble proteins and IAA (IAA, IAA-Ala, IAA-Trp, IAA-Val-Me) and CK (tZOG, K), GA (GA19) *(0*.*01<p<0*.*05)*, the intermediates produced by sucrose in photosynthesis can be used as raw materials for the synthesis of phytohormones, which are involved in the synthesis of Auxin (IAA, IAA-Ala, TRP, IAA-Trp), CK (KR, tZR, K, mT9G), GA19, and JA (JA-ILE), and the synthesis of soluble starch, IAA (IAA, IAA-Val-Me), and GA (GA19). Ala, IAA-Val-Me, IAA-Trp), CK (tZOG, KR, BAP9G, DHZR, mT9G, K), and GA (GA19) were significantly positively correlated *(0*.*01<p<0*.*05)*, and the hormones provided carbon skeleton and energy by regulating the physiological state of the plant, which induced the plant to convert more photosynthesis products into sugars and transported them to the bud site Accelerating the initiation of flower bud differentiation. Cytokinin showed significant positive correlation *(0*.*01<p<0*.*05)* with sucrose, soluble starch and malondialdehyde, which is conducive to the promotion of flower differentiation, and it is closely related to the accumulation of carbohydrates, cytokinin affects the activity of key enzymes such as sucrose synthase by regulating the expression of related genes, so that sucrose and soluble starch accumulate in the buds, and malondialdehyde content is increased to promote resistance to adversity, and provide the provide energy and material basis for flower bud differentiation (Fig 6-A, C).

**Fig 6.**
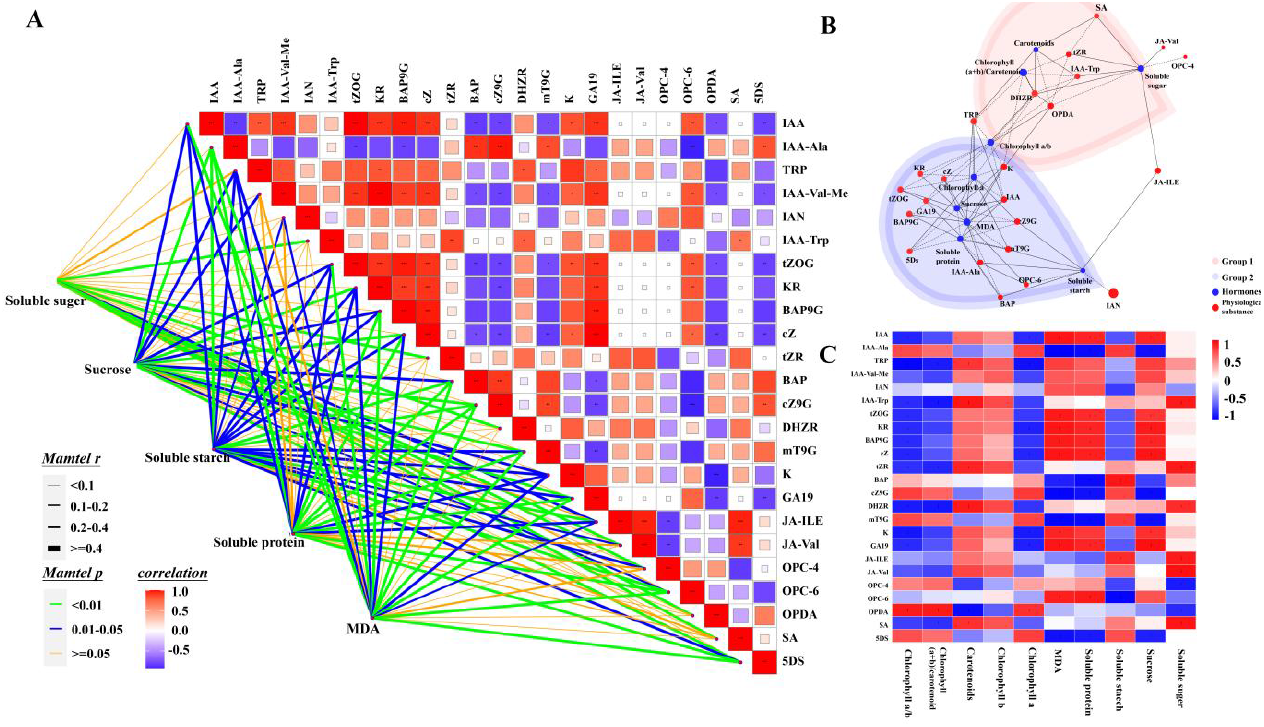
Correlation network of physiological and hormonal substances for flower bud differentiation Note:A: Manteltest analysis; B: Correlation network diagram; C: Clustering heat map.

## 7 Discussion

The transition from nutrient growth to reproductive growth in plants is regulated through a complex network of interactions between genetic factors and external environmental conditions. After sensing changes in the external environment, plants adjust the bud differentiation process to adapt to the changing environmental conditions through the transmission and integration of physiological substances and hormone signalling networks. Exploring the morphological and physiological characteristics of flower bud differentiation in Bombax Bombax ceiba is of great significance in revealing the growth and development of Bombax Bombax ceiba, precisely regulating the flowering period in agricultural production, improving the yield and quality of the crop, and cultivating rare varieties in the field of horticulture.

*Bombax ceiba* flower buds are mixed buds of monoecious and dioecious plants, and according to the time of germination, they can be divided into winter and spring differentiation type, which is formed by the differentiation of terminal buds through the induction of flower formation. The transition from nutrient growth to reproductive growth in *Bombax ceiba* flower bud differentiation is divided into two stages, one is the transformation from the growing point to the flower bud known as floral primordium development (from late October to January of the following year, which lasts 60 d), and the other is the maturation of the floral organ (from early January to March of the following year, which lasts 75 d). This is similar to the results of Hu Shuangling and Zhu (Hu Shuangling and Lin Gongtian, 2024; Zhu and Zhang Fangqiu, 2021). Physiological substances are essential energy sources for maintaining normal life activities of flowers (Lin, B., & Wang, D, 2023). The results show that the same plant is affected by the photoperiod and temperature, the plant physiological rhythm of its own growth process there are differences, which in turn affects the accumulation of physiological substances, the early soluble sugar and soluble starch content increased hydrolysis into directly usable energy substances for the initiation of flower bud differentiation, the large amount of sugar in the leaves, phloem, and buds to transport nutrients for the development of the pollen mother cells in the late stage, and then with the warming of the temperature gradually slowed down the stress. Soluble proteins are involved in the construction of cell membrane morphology, early flowering *Bombax ceiba* soluble protein content increases through a large number of enzymes to catalyse a variety of biosynthetic reactions, such as nucleic acid synthesis, hormone synthesis, etc., to ensure that the cells can carry out the metabolic activities smoothly, laying the groundwork for the initiation of the buds and the development of the foundation. Malondialdehyde can play a certain osmotic regulation of water in the plant body under adverse conditions. When the temperature gradually close to 24 °C to reach the appropriate growth temperature of *Bombax ceiba*, with the temperature rising osmotic stress continues to increase, the buds appeared in varying degrees of physiological water shortage, prompting the buds continue to produce and accumulate malondialdehyde in the body. When the temperature exceeded 28°C, the increase in external temperature slowed down, and the change in membrane peroxides slowed down accordingly (Hiroyuki Tsuji, & Moeko Sato, 2024).

Chlorophyll provides energy and material basis for plants through photosynthesis. Previous studies have rarely measured the trends of chlorophyll and carotenoids during bud differentiation, and further studies are needed to investigate the regulatory mechanisms of chlorophyll and carotenoids in the bud differentiation process of *Bombax ceiba*. Chlorophyll a absorbs and converts light energy to convert carbon dioxide and water into organic matter, and chlorophyll b assists chlorophyll a in photosynthesis. It is worth noting that in this study, the trends of changes in chlorophyll a and chlorophyll b in *Bombax ceiba* were found to be opposite at the stage of bud differentiation, and it is speculated that the photosynthetic efficiency is improved by adjusting its own metabolic pathway to reduce the input of chlorophyll synthesis during the physiological differentiation stage. Carotenoids, as precursors of abscisic acid and solanum lactone, act synergistically with light signalling to regulate bud differentiation. In past studies, epigenetic processes have been associated with carotenoid biosynthesis, accumulation and degradation in plant growth and development, and a gradual decrease in carotenoid content at the stage of bud differentiation has been speculated to be associated with a feedback effect with other genes in the carotenoid biosynthesis pathway as a way of regulate carotenoid biosynthesis in plants.

Plant hormones are the most important endogenous signalling players. Studies have shown that there are synergistic and interacting resistance among hormones. By determining the changes in hormone content during flower bud differentiation in Bombax ceiba, it was found that ABA (ABA, ABA-GE), as a hormone that inhibits growth and promotes dormancy and senescence, changed significantly during phase I and phase II. Differential metabolites related to hormone regulation of bud differentiation were mainly concentrated in metabolism and environmental information processing. *Bombax ceiba* regulated hormone concentration by changing the expression of different kinds of endogenous hormone synthesis and metabolism genes during bud differentiation, and phytohormones in turn regulated *Bombax ceiba* bud differentiation by causing the expression of downstream genes through the signal transduction pathway. The up-regulated expression was mainly concentrated in Auxin, CK, GA and SA, and the down-regulated expression was mainly concentrated in Auxin, CK, JA and 5Ds. The roles of CK and Auxin were the most significant in the process of *Bombax ceiba* bud differentiation, and the metabolic pathways involved in the early biological process as a response to the stimulus, and transmembrane transporter, cleaving enzyme activity, and structural molecule activity were related to them. The co-expression network reveals that physiological substances and endogenous hormones interact with each other. Hormones regulate carbohydrate metabolism and partitioning, and physiological substances in turn regulate hormone synthesis and signalling in a feedback manner. Under low-temperature stress at the early stage of bud differentiation, the physiological substances MDA, sucrose, soluble protein, and chlorophyll a were positively correlated, and at this time, a large number of reactive oxygen radicals were generated in the plant, which attacked unsaturated fatty acids in the cell membrane and triggered lipid peroxidation to produce malondialdehyde.The Mantel test analysis and the clustering heatmap indicated that sucrose Sucrose was involved in the synthesis of growth hormone and cytokinins, and it induced the transport of growth hormone. carrier genes (CmPIN1 and CmAUX1) and inhibits the expression of growth hormone signalling genes (CmIAA3, CmIAA14 and CmIAA16), and down-regulates the expression of growth hormone response factor gene CmARF2 in the terminal buds, which enhances growth hormone transport and reduces the level of endogenous growth hormone in the terminal buds, thus promoting the growth of flower buds. Soluble proteins showed a significant positive correlation with Auxin and GA. Soluble proteins and Auxin showed a positive correlation, and soluble proteins such as Klotho proteins affected the expression of genes related to renal tubular epithelial cell fibrosis by inhibiting the STAT3 phosphorylation pathway in a high glucose environment. Meanwhile, PIN proteins, the growth hormone transport carrier, interacted with other soluble proteins, and played a polar role in the polar transport of growth hormone The growth hormone transport carrier PIN protein interacts with other soluble proteins and plays a key role in polar transport of growth hormone.

In summary, the network of phytohormones and physiological substances regulating bud differentiation is a highly complex and ordered system, in which various phytohormones act in time and space in a balanced manner, controlling nutrient metabolism, regulating physiological processes at the cellular level to the regulation of gene expression, and comprehensively and finely regulating the whole process of bud differentiation from its initiation to its completion, in order to adapt to the various environmental changes in the process of plant growth and development. The following is a list of some of the most important aspects of the plant’s development.

## 8 Acknowledgements

We are also grateful to the Key Laboratory of Genetics and Germplasm Innovation of Tropical Speciality Forestry and Floriculture of the Ministry of Education for its strong support of plant physiological experiments.

## 11 Data and resource availability

All relevant data and resource can be found within the article and its supplementary information.

## 12 Diversity and inclusion statement

No Diversity and Inclusion Statement.

